# Manganese Restricts HIV-1 Infection by Suppressing Viral Reverse Transcription in Human Macrophages

**DOI:** 10.64898/2026.11.27.769786

**Authors:** Hongyan Sui, Rosana Wiscovitch-Russo, Whitney Bruchey, Jordan Metz, Adam Rupert, Sylvain Laverdure, Tomozumi Imamichi

## Abstract

Despite the clinical success of combination antiretroviral therapy, the persistence of latent viral reservoirs, cumulative drug toxicities, and emerging viral resistance necessitate novel host-directed strategies against human immunodeficiency virus type 1 (HIV-1). Manganese (Mn) has recently emerged as an essential regulator of innate antiviral immunity, yet its activity and mechanism of action across primary human HIV-1 target cells remain poorly defined. In this study, we evaluated the antiviral effects of Mn in primary human monocyte-derived macrophages, dendritic cells, and CD4+ T cells following pretreatment or post-treatment relative to HIV-1 challenge. We found that Mn inhibited HIV-1 infection in primary human macrophages in a dose-dependent manner, achieving up to 80% and 90% suppression following pretreatment and post-treatment at 20 µM or 25 µM, respectively, with no apparent changes in cell morphology. Strikingly, this antiviral activity was cell-type-specific, occurring in macrophages but not in primary dendritic cells or CD4+ T cells. Mechanistic mapping of the viral replication cycle revealed that Mn suppressed intracellular HIV-1 reverse transcription; however, cell-free enzymatic assays confirmed that Mn at antiviral concentrations did not directly inhibit viral reverse transcriptase catalytic activity. Furthermore, mRNA profiling of 10 selected transcripts demonstrated that Mn induced sustained expression of multiple host genes associated with innate antiviral defense and immune regulation. Together, these findings identify reverse transcription as a primary checkpoint of Mn-mediated HIV-1 restriction in human macrophages and suggest that Mn promotes a host-directed antiviral state rather than directly blocking viral enzymes, supporting further investigation of metalloimmunological pathways for anti-HIV-1 interventions.

## Introduction

Although the advent of combination antiretroviral therapy (ART) has dramatically reduced HIV-related morbidity and mortality, HIV-1 infection remains a major global public health challenge [1, 2]. Current ART regimens effectively suppress active viral replication to undetectable levels in peripheral blood, yet they do not eradicate integrated proviral DNA from long-lived cellular and anatomical reservoirs, such as resting memory CD4+ T cells and tissue macrophages [3, 4]. Consequently, interruption of lifelong treatment almost invariably leads to rapid viral rebound [5, 6]. Moreover, sustained adherence to daily regimens is complicated by pill fatigue, cumulative drug toxicities, and chronic immune activation [7].

A central target of current antiretroviral regimens is reverse transcriptase (RT), a conserved enzyme encoded across retroviruses such as HIV-1 and other retroelements [8, 9]. Functioning as a specialized DNA polymerase, RT utilizes both RNA and DNA templates to catalyze two essential steps in viral replication: synthesizing minus-strand DNA from the viral RNA genome and subsequently generating plus-strand DNA [10, 11]. Because these activities are indispensable for viral propagation, nucleoside and non-nucleoside RT inhibitors remain cornerstones of modern ART [12]. However, the high mutation rate of HIV-1 RT drives the ongoing emergence of drug-resistant viral variants, underscoring the continued need for novel therapeutic strategies that suppress viral replication through alternative, host-directed mechanisms [13].

Transition metals have recently emerged as critical regulators of host– pathogen interactions and innate immunity [14–16]. Among essential trace elements, manganese (Mn) functions not only as an indispensable cofactor for cellular metabolic and antioxidant enzymes, but also as a potent endogenous alarm signal during viral infection [17, 18]. Upon viral entry, Mn is mobilized from membrane-bound organelles, such as the mitochondria and Golgi apparatus, into the cytosol [19]. Once released, cytosolic Mn markedly increases the sensitivity of the DNA sensor cyclic GMP-AMP synthase (cGAS) to double-stranded DNA and enhances cGAMP– STING binding affinity, thereby lowering the activation threshold for type I interferon (IFN) and pro-inflammatory cytokine production [19, 20]. Beyond canonical cGAS–STING sensing, accumulating evidence including our previous studies demonstrates that Mn exerts broad-spectrum antiviral activity against both DNA and RNA viruses via cGAS–STING-independent signaling cascades, including the phosphorylation of ATM and TBK1 and the broad induction of interferon-stimulated genes (ISGs) [21–23]. Accordingly, Mn treatment has been shown to suppress the replication of diverse DNA and RNA viruses, including herpes simplex virus 1 (HSV-1) [19, 22], hepatitis B virus (HBV) [24, 25], influenza virus [26], and foot-and-mouth disease virus (FMDV) [27].

Despite these major insights into Mn-mediated antiviral immunity, the therapeutic potential and precise mechanisms of Mn against HIV-1 infection in primary human target cells remain incompletely defined. As a retrovirus, HIV-1 generates double-stranded DNA intermediates during reverse transcription that are susceptible to cytosolic DNA sensing, yet the virus employs multiple evasion strategies to subvert host innate recognition [28, 29]. Whether physiological or sub-toxic Mn concentrations restrict HIV-1 across distinct primary immune cell subsets, and whether such restriction occurs through direct enzymatic blockade or host-mediated innate pathways remains largely unknown. So, in this study, we investigated the anti-HIV-1 activity of Mn and sought to define the underlying molecular mechanisms driving viral suppression. The findings will provide new insights into the metalloimmunological control of retroviral infection and supporting future host-directed anti-HIV-1 strategies.

## Material and Methods

### Isolation and Culture of Primary Human Cells

Peripheral blood mononuclear cells (PBMCs) were isolated from healthy donor leukopacks (NIH Blood Bank, Bethesda, MD, USA). CD14^+^ monocytes were purified using CD14 MicroBeads (Miltenyi Biotec). To generate monocyte-derived macrophages (MDMs), CD14^+^ monocytes (1 × 10^7^ cells per 10-cm dish) were differentiated for 7 days in macrophage serum-free medium (Thermo Fisher Scientific) supplemented with 25 ng/mL M-CSF (R&D Systems) and subsequently maintained in D10 medium (DMEM with 10% FBS). CD4^+^ T cells were isolated from the CD14^−^PBMC fraction using CD4 MicroBeads (Miltenyi Biotec) and activated with 5 µg/mL phytohemagglutinin (PHA; Sigma-Aldrich) for 3 days in RPMI 1640 supplemented with 10% FBS, 25 mM HEPES, and 5 µg/mL gentamicin. Immature dendritic cells (iDCs) were differentiated by culturing CD14^+^ monocytes (0.5 × 10^6^ cells/mL) for 7 days in RPMI 1640 containing 10% FBS, 2 mM L-glutamine, 25 mM HEPES, 5 µg/mL gentamicin, 50 ng/mL GM-CSF, and 50 ng/mL IL-4 (R&D Systems), with half-medium replenishment every 3–4 days. All cells were maintained at 37°C with 5% CO_2_.

### RNA Extraction and Quantitative Real-Time RT-PCR

Total RNA was extracted using the RNeasy Mini Kit (QIAGEN) and reverse-transcribed using TaqMan Reverse Transcription Reagents with random hexamers (Thermo Fisher Scientific). Quantitative PCR (qPCR) was performed using TaqMan Gene Expression Assays on a QuantStudio 7 Pro Real-Time PCR System (Thermo Fisher Scientific) for 40 cycles (95°C for 15 s; 60°C for 1 min). Relative expression was calculated via the 2^−^ΔΔCt method using *GAPDH* as the endogenous control and normalized to untreated controls.

### Preparation of HIV-1 Virus Stocks

Replication-competent HIV-1 AD8 was produced by transfecting HEK293T cells with pAD8 [30]. VSV-G-pseudotyped luciferase reporter virus (HIV-Luc-V) was generated by cotransfecting HEK293T cells with pHIV-1NL4/3-Luciferase-ΔEnv and pVSV-G [31–33]. Vpr–β-lactamase-containing particles (HIV-1 Vpr-BlaM) were produced by cotransfecting HEK293T cells with pHIV-1NL4.3-ΔEnv-ΔVpr, pVpr-BlaM, pVSV-G, and pAdvantage [34]. Transfections were performed in DMEM using TransIT-293 reagent (Mirus Bio, Houston, TX, USA), as previously described [35]. Culture supernatants were concentrated by ultracentrifugation (100,000 × g, 2 h, 4°C) through a 20% sucrose cushion (10 mM HEPES, 150 mM NaCl) and resuspended in D10. Viral p24 levels were quantified by HIV-1 p24 ELISA (PerkinElmer, Boston, MA, USA), and infectious titers (TCID_50_) were determined by endpoint dilution.

### HIV-1 Infection and Replication Assays

For single-round infection [36], MDMs (3 × 10^5^ cells/well in 24-well plates) were inoculated with HIV-Luc-V for 2 h at 37°C, washed three times with prewarmed D10, and cultured for 48–72 h. Cells were lysed in 1× Passive Lysis Buffer, and luciferase activity was measured using the Luciferase Assay System (Promega, Madison, WI, USA) on a Tecan plate reader (Morrisville, NC, USA). For spreading replication, MDMs (3 × 10^5^ cells/well) were infected with HIV-1 AD8 (5,000 TCID_50_ per 10^6^ cells) for 2 h at 37°C under continuous rotation, washed three times with D10, and cultured for 10 days with half-medium replacement (Mn-free D10) every 3–4 days. Supernatant p24 was quantified by ELISA (Revvity) [35].

### HIV-1 Viral Binding Assay

MDMs (1 × 10^6^ cells/well in 6-well plates) were incubated with HIV-Luc-V (500 ng p24/mL) or HIV-1 AD8 (5,000 TCID_50_ per 10^6^ cells) for 2 h at 4°C, washed with ice-cold PBS, and lysed in 100 µL RIPA buffer (Boston BioProducts, Milford, MA, USA) containing protease and phosphatase inhibitors (Thermo Fisher Scientific) for 15 min at 4°C. After centrifugation (3,500 rpm, 10 min, 4°C), cell-associated p24 was measured by ELISA and normalized to total protein determined via BCA assay (Thermo Fisher Scientific).

### HIV-1 Fusion Assay and Imaging Flow Cytometry

Viral fusion was measured using the Vpr-BlaM assay [37]. MDMs (3 × 10^6^ cells) were inoculated with HIV-1 Vpr-BlaM (500 ng p24/mL) for 2 h at 37°C, washed three times, loaded with CCF2-AM (Thermo Fisher Scientific), and incubated in CO_2_-independent medium for 1 h at 37°C in the dark. Cells were detached with Accutase and analyzed on an ImageStream X Mark II Imaging Flow Cytometer (Amnis, Cytek Biosciences, Seattle, WA, USA; 40× objective, low flow rate, high sensitivity). CCF2 was excited at 405 nm (10 mW), with cleaved (blue) and uncleaved (green) fluorescence detected in channels 7 and 8, respectively; side scatter (785 nm, 2.66 mW) was collected in channel 12, and brightfield images in channels 1 and 9 (∼50,000 events/sample). Using IDEAS software (v6.4), in-focus single cells were gated by brightfield Gradient RMS (>50), area (200–600), and aspect ratio (>0.6).

### Flow Cytometry Analysis

MDMs (1 × 10^6^ cells/well in 6-well plates) were treated with 25 µM Mn or left untreated in D10 for 48 h, washed with DPBS, and detached using Accutase. Cells were washed in DPBS (500 × g, 5 min), blocked with Fc Receptor Blocker (Innovex Biosciences, Richmond, CA, USA) for 30 min at room temperature in the dark, and washed twice in staining buffer (DPBS containing 2% BSA and 0.5% NaN_3_). Cells were then stained for 15 min at room temperature in the dark with BV421-conjugated anti-CD4 (BioLegend, San Diego, CA, USA; Cat# 317434) or BV421 isotype control (BioLegend; Cat# 400158), and APC-Cy7-conjugated anti-CCR5 (BD Biosciences, Franklin Lakes, NJ, USA; Cat# 557755) or APC-Cy7 isotype control (BioLegend; Cat# 400128). After three washes in staining buffer, cells were acquired on an LSRFortessa flow cytometer (BD Biosciences) and analyzed using FCS Express v7 (DeNovo Software, Pasadena, CA, USA).

### Quantification of HIV-1 Late Reverse Transcription Products

Total cellular DNA was extracted from infected MDMs using the QIAamp DNA Mini Kit (QIAGEN; Cat# 51304). HIV-1 late reverse transcription (RT) products were quantified by qPCR or digital PCR (dPCR). For qPCR, reactions were performed using TaqMan Universal PCR Master Mix (Thermo Fisher Scientific; Cat# 4304437) on a QuantStudio 7 Pro System (50°C for 2 min, 95°C for 10 min, followed by 40 cycles of 95°C for 15 s and 60°C for 1 min) using primers MH531 (5′-TGTGTGCCCGTCTGTTGTGT-3′), MH532 (5′-GAGTCCTGCGTCGAGAGAGC-3′), and probe LRT-P (5′-FAM-CAGTGGCGCCCGAACAGGGA-TAMRA-3′), as previously described[38]. However, to amplify HIV-Luc-V, the reverse primer MH532 was modified to 5’-GAGTCCTGCGTCGAGAGATC-3’ via a G-to-T substitution in nucleotide 19. Cellular RNase P was quantified in parallel using TaqMan RNase P Detection Reagents (Thermo Fisher Scientific; Cat# 4316831). Copy numbers were calculated from plasmid DNA standard curves (10 to 1 × 10^6^ copies). For dPCR, absolute copy numbers of HIV-1 late RT products and RNase P were quantified in triplicate on a QIAcuity Eight Digital PCR System (QIAGEN) without standard curves. For both methods, late RT levels were normalized to RNase P and expressed as copies per 1,000 cells.

### Cell-Free HIV-1 Reverse Transcriptase Activity Assay

HIV-1 RT activity was measured using a fluorescence-based assay kit (ProFoldin, Hudson, MA, USA; Cat# HIV100KE) in 96-well plates (60 µL/well). Reaction mixtures containing 1× assay buffer, synthetic template– primer hybrid, dNTPs, and recombinant HIV-1 RT were supplemented with MnCl_2_ (25 µM, 50 µM, or 0–10 mM) or an equivalent volume of sterile water (0 µM control), alongside no-template controls (NTC). After incubation for 60 min at 37°C, reactions were terminated with 30 µL of 1× fluorescent dye in 50 mM EDTA. Fluorescence intensity (excitation 485 nm, emission 535 nm) was measured on a Tecan Spark microplate reader (Morrisville, NC, USA), and NTC background values were subtracted.

### MultiScribe Reverse Transcriptase Activity Assay

To evaluate the effect of Mn on MultiScribe reverse transcriptase (RT, Thermo Fisher Scientific), cDNA synthesis reactions were performed using TaqMan Reverse Transcription Reagents with random hexamers (Thermo Fisher Scientific) and total RNA isolated from untreated MDMs (RNA input: 500 ng). Reactions were prepared in standard reaction buffer containing 5.5 mM MgCl_2_ supplemented with increasing concentrations of MnCl_2_ (0–5.0 mM), or in buffer in which 5.5 mM MgCl_2_ was completely replaced with 5.5 mM MnCl_2_. Following reverse transcription, cDNA copy numbers of *GAPDH* and *CDKN1A* were quantified using TaqMan Gene Expression Assays on a QIAcuity Eight Digital PCR System (QIAGEN). Enzymatic activity was normalized to the standard reaction containing 5.5 mM MgCl_2_ alone and expressed as a percentage of the control.

### siRNA Transfection

MDMs were reverse-transfected with 25 nM ATM-targeting siRNA (siATM; Horizon Discovery; Cat# L-003201-00-0005) or non-targeting control siRNA (siCtrl; Horizon Discovery; Cat# D-001810-10-05) using Lipofectamine RNAiMAX (Thermo Fisher Scientific) and incubated for 48 h at 37°C with 5% CO_2_ prior to Mn treatment and HIV-1 infection.

### Western Blot Analysis

MDMs treated with 25 µM Mn for the indicated durations were washed with ice-cold PBS and lysed in RIPA buffer (Boston BioProducts) supplemented with protease and phosphatase inhibitor cocktails (Thermo Fisher Scientific. Protein concentrations were determined using a BCA protein assay kit (Thermo Fisher Scientific). Equal amounts of protein (20 µg per lane) were resolved by SDS–PAGE, transferred onto nitrocellulose membranes, and probed overnight at 4°C with primary antibodies against phosphorylated SAMHD1 (phospho-SAMHD1 Thr592; Cell Signaling, Cat# 89930), total SAMHD1 (Cell Signaling, Cat# 12361), and β-actin (Sigma-Aldrich, Cat# A5316), followed by HRP-conjugated secondary antibodies (GE Healthcare, Chicago, IL). Protein bands were visualized with Amersham ECL Prime Western Blotting Detection Reagent (Cytiva, Cat# RPN2236) prior to visualizing the protein bands on Chemiluminescent Western Blot Imager Azure 300 (Azure Biosystems, Dublin, CA).

### Statistical Analysis

Unless otherwise indicated, data were obtained from at least three independent experiments and are presented as mean ± standard deviation (SD). Statistical significance was determined using an unpaired Student’s t-test or one-way or two-way analysis of variance (ANOVA) with appropriate post hoc multiple-comparison tests. A p-value < 0.05 was considered statistically significant (* p < 0.05, ** p < 0.01, *** p < 0.001, **** p < 0.0001).

## Results

### Mn dose-dependently suppresses HIV-1 infection in human primary macrophages

To evaluate the antiviral activity of Mn against HIV-1 infection, human monocyte-derived macrophages (MDMs) were pretreated with increasing concentrations of MnCl_2_ for 24 or 48 h before infection with VSV-G-pseudotyped HIV-1 expressing luciferase (HIV-Luc-V). Mn treatment inhibited single-round HIV-1 infection in a dose-dependent manner, with 48 h of pretreatment producing greater inhibition than 24 h. Notably, Mn at a concentration of 20 µM suppressed viral infection by more than 80% following 48 h of pretreatment (Figure 1A). No overt changes in cell morphology were observed by microscopy at the concentrations tested.

**Fig 1.**
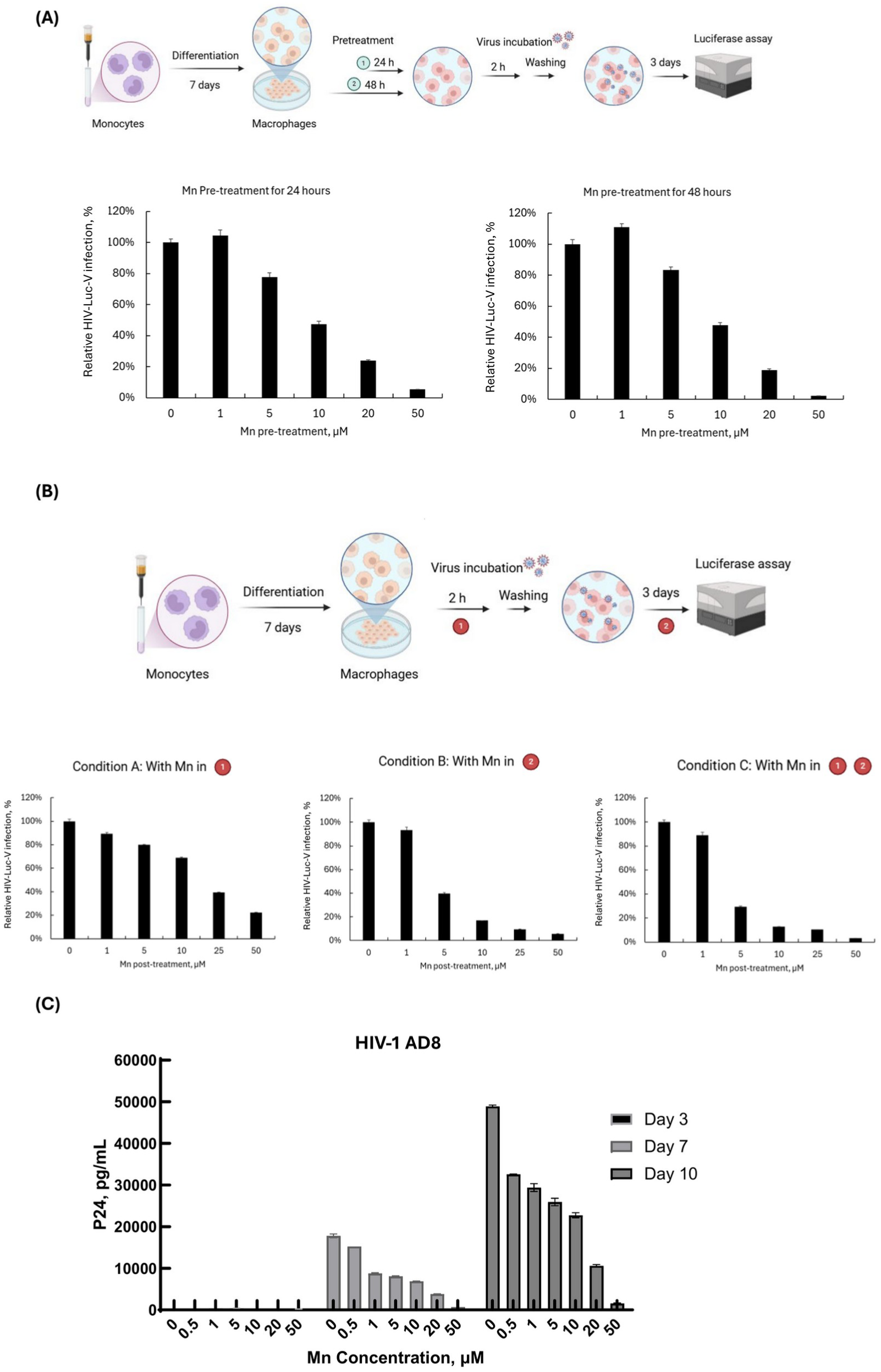
Manganese (Mn) treatment dose-dependently inhibits HIV-1 replication in primary human macrophages. Primary human monocyte-derived macrophages (MDMs) were seeded in 24-well plates and either pretreated or post-treated with Mn at the indicated doses. Viral replication in infected cells was quantified via luciferase activity assays or p24 ELISA. **(A)** Schematic diagram of the pretreatment experimental design (Created in BioRender. *Sui, H. (2027)* https://BioRender.com/h4w75d3) and corresponding results, showing that pretreatment with Mn for 24 h or 48 h results in a dose-dependent inhibition of HIV-Luc-V infection. **(B)** Schematic diagram of the post-treatment experimental design (*Created in BioRender. Sui, H. (2027)* https://BioRender.com/h4w75d3) and corresponding results, showing dose-dependent inhibition of HIV-Luc-V. Mn was introduced during viral inoculation (Condition A), after viral washout (Condition B), or maintained during both inoculation and post-infection incubation (Condition C). **(C)** Pretreatment with Mn demonstrates dose-dependent inhibition of wild-type (WT) HIV-1 AD8 infection. Cell supernatants were collected at 3, 7, and 10 days post-infection (dpi). Data are presented as mean ± SD (n = 3 technical replicates) and are representative of three independent donors.

We next investigated whether Mn could also inhibit HIV-1 infection when administered during or after viral inoculation. MDMs were treated with increasing concentrations of MnCl_2_ under three conditions: Mn exposure during the 2-h viral inoculation period only (condition A), Mn addition after viral inoculation and maintenance in the culture medium (condition B), or Mn exposure during both viral inoculation and subsequent culture (condition C). As shown in Figure 1B, Mn treatment under all three conditions suppressed HIV-1 infection, with greater inhibition observed when Mn was maintained in the culture medium following viral inoculation compared with exposure during inoculation alone.

To determine whether the antiviral effect of Mn extended to replication-competent HIV-1, MDMs were pretreated with increasing concentrations of MnCl_2_ before infection with the macrophage-tropic HIV-1 strain AD8 (HIV-1 AD8). As shown in Figure 1C, Mn pretreatment reduced HIV-1 p24 production in a dose-dependent manner, with the inhibitory effect sustained for up to 10 days post-infection.

Given the potent antiviral activity of Mn observed in primary human macrophages, we next investigated whether similar Mn pretreatment could inhibit HIV-1 infection in other primary immune cell types, including CD4+ T cells and monocyte-derived immature dendritic cells (iDCs), both of which play important roles in HIV-1 pathogenesis. As shown in Supplementary Figure S1, Mn pretreatment did not significantly inhibit HIV-VSV-G-Luc infection in either CD4+ T cells (Figure S1A) or iDCs (Figure S1B). These findings suggest that the antiviral activity of Mn is cell-type dependent, with a preferential inhibitory effect observed in primary macrophages under the conditions tested.

Collectively, these results demonstrate that Mn potently suppresses HIV-1 infection in primary human macrophages, with antiviral activity against both single-round and replication-competent HIV-1. Furthermore, the enhanced inhibition observed following prolonged Mn exposure suggests that treatment duration is an important determinant of its antiviral efficacy.

### Mn pretreatment does not substantially block HIV-1 binding or viral fusion in human primary macrophages

To identify the stage of the HIV-1 replication cycle affected by Mn treatment, we first investigated whether Mn interferes with viral attachment to macrophages. MDMs were pretreated with 25 µM MnCl_2_ for 48 h before exposure to either VSV-G-pseudotyped HIV-1 (HIV-Luc-V) or replication-competent HIV-1 AD8. Mn pretreatment did not decrease, and in fact significantly increased HIV-Luc-V binding, whereas a modest reduction of approximately 20% was observed for HIV-1 AD8 binding (Figure 2A).

**Fig 2.**
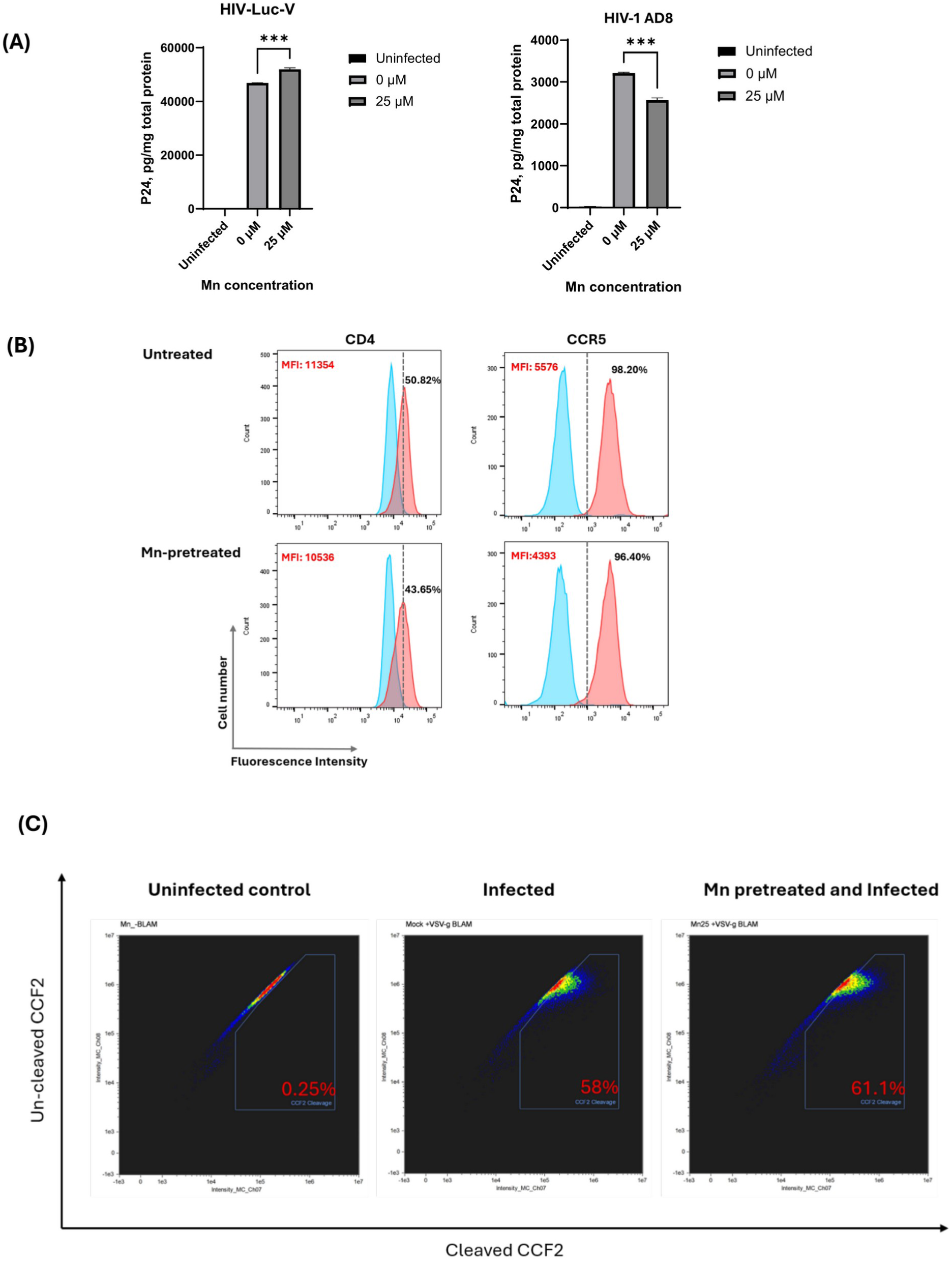
Mn pretreatment does not substantially impair HIV-1 binding or VSV-G-mediated viral fusion in primary human macrophages. **(A)** MDMs were seeded in 24-well plates and pretreated with 25 µM MnCl_2_ for 48 h. Cells were subsequently infected with HIV-Luc-V or HIV-1 AD8 at 4°C for 2 h to allow viral binding. Following incubation, cells were thoroughly washed to remove unbound viral particles. Whole-cell lysates were collected, and cell-associated viral p24 levels were quantified by ELISA. Data is presented as mean ± SD (n = 3) and is representative of at least three independent donors. Statistical significance was determined by one-way ANOVA with Dunnett’s multiple comparisons test: *** p < 0.001. **(B)** Cell-surface expression of CD4 and CCR5 in untreated (top panels) and MnCl_2_-pretreated (bottom panels) MDMs was assessed by flow cytometry. Blue histograms represent isotype controls, whereas red histograms indicate CD4- or CCR5-specific antibody staining. The x- and y-axes represent fluorescence intensity and cell count, respectively. Data are representative of three independent experiments with similar results. **(C)** MDMs were pretreated with 25 µM MnCl_2_ for 48 h and subsequently either mock-infected or inoculated with VSV-G-pseudotyped HIV-1 particles containing Vpr–β-lactamase (Vpr-BlaM) for 1 h at 37°C. Following viral incubation, cells were loaded with the fluorescent substrate CCF2-AM, and viral fusion was assessed using the BlaM fusion assay. Mock-uninfected cells were used as negative controls to establish background gating. Representative flow cytometry dot plots from three independent experiments are shown.

We next examined whether Mn treatment altered the surface expression of CD4 and CCR5, the primary receptor and coreceptor required for HIV-1 AD8 entry. Flow cytometric analysis revealed a modest reduction in the percentage of CD4-positive macrophages following Mn pretreatment (50.82% in untreated cells versus 43.65% in Mn-treated cells), accompanied by a slight decrease in mean fluorescence intensity (MFI; 11,354 versus 10,536). Although the percentage of CCR5-positive cells remained largely unchanged (98.20% versus 96.40%), CCR5 MFI decreased from 5,576 to 4,393 following Mn treatment (Figure 2B). These observations suggest that Mn pretreatment modestly alters CD4 and CCR5 surface expression, which may contribute to the slight reduction in HIV-1 AD8 binding. However, these changes are unlikely to fully account for the pronounced inhibition of HIV-1 infection observed following Mn treatment.

To further investigate whether Mn affects viral membrane fusion, we performed a fusion assay using VSV-G-pseudotyped HIV-1 carrying Vpr–β-lactamase (HIV-VSV-G-BlaM). Given the modest reduction in HIV-1 AD8 binding following Mn pretreatment, VSV-G-pseudotyped HIV-1 was specifically selected to minimize potential interference from CD4/CCR5-dependent viral attachment, allowing us to evaluate VSV-G-mediated fusion independently of these receptors. Viral fusion was quantified by measuring CCF2 substrate cleavage using imaging flow cytometry. The percentage of cells exhibiting CCF2 cleavage was comparable between untreated and Mn-treated macrophages (58.0% and 61.1%, respectively), indicating that Mn pretreatment did not inhibit VSV-G-mediated viral fusion (Figure 2C).

Overall, these findings demonstrate that Mn pretreatment does not substantially impair HIV-1 binding or VSV-G-mediated viral fusion in human primary macrophages, despite modest changes in CD4 and CCR5 surface expression. These observations suggest that the potent antiviral activity of Mn is unlikely to be primarily mediated by inhibition of viral attachment or entry, supporting a mechanism of restriction at a post-entry stage of the HIV-1 replication cycle.

### Mn potently inhibits reverse transcription step but not through directly suppressing HIV-1 reverse transcriptase enzyme

Having established that Mn pretreatment does not substantially impair viral binding or VSV-G-mediated fusion, we next investigated whether Mn restricts HIV-1 infection at the reverse transcription stage. To minimize potential interference from CD4/CCR5-dependent viral entry, we used VSV-G-pseudotyped HIV-1 to evaluate the effect of Mn on viral reverse transcription. HIV-1 late reverse transcription (RT) products were quantified by real-time PCR and normalized to cellular RNase P copy numbers. As shown in Figure 3A, Mn pretreatment markedly reduced the accumulation of HIV-1 late RT products, indicating that Mn suppresses viral reverse transcription at either 25 µM or 50 µM.

**Fig 3.**
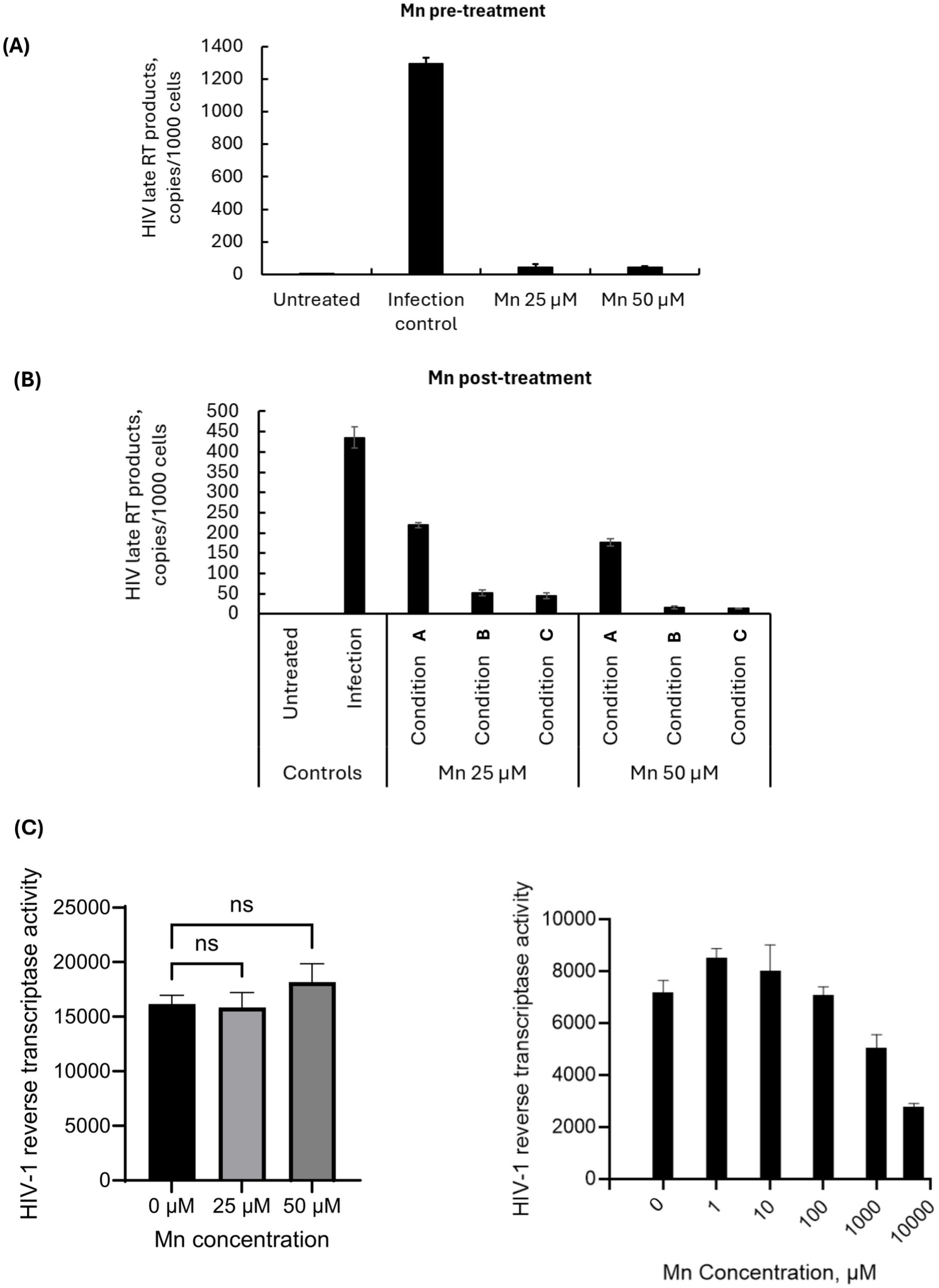
Mn predominantly suppresses HIV-1 late reverse transcription products following pretreatment or treatment during infection. **(A)** Effect of Mn pretreatment on HIV-1 reverse transcription. MDMs were seeded in 24-well plates and pretreated with 25 or 50 µM MnCl_2_ for 48 h. Cells were subsequently inoculated with HIV-Luc-V for 2 h at 37°C. Following inoculation, cells were thoroughly washed to remove unbound viral particles. Total cellular DNA was extracted at 48 h post-infection, and HIV-1 late reverse transcription (RT) products were quantified by quantitative real-time PCR (qPCR). HIV-1 late RT copy numbers were normalized to cellular RNase P copy numbers. Data are shown as mean ± SD (n = 3) copies per 1,000 cells and are representative of at least three independent donors. **(B)** Effect of Mn treatment timing on HIV-1 late reverse transcription. MnCl_2_ was administered under three conditions, as illustrated in Fig 1B: Condition A, during the 2-h viral inoculation only; Condition B, after viral inoculation; and Condition C, during both inoculation and the post-inoculation period. At 48 h post-infection, total cellular DNA was extracted, and HIV-1 late RT products were quantified by qPCR and normalized to RNase P copy numbers as described in panel A. **(C, D)** Mn does not directly inhibit HIV-1 reverse transcriptase (RT) enzymatic activity at antiviral concentrations. The effects of Mn on HIV-1 RT activity were evaluated using a cell-free HIV-1 Reverse Transcriptase Assay Kit (ProFoldin) according to the manufacturer’s instructions. MnCl_2_ was tested at final concentrations of 25 and 50 µM (C) or across a broader concentration range of 0–10 mM (D). HIV-1 RT enzymatic activity was measured to assess the concentration-dependent effects of Mn (n = 3).

We further examined the effect of Mn on HIV-1 reverse transcription under different treatment conditions during posttreatment, consistent with those used in Figure 1B. Mn was administered during viral inoculation, after inoculation, or during both periods. As shown in Figure 3B, Mn treatment reduced the accumulation of late RT products normalized to RNase P copy numbers, with greater inhibition observed when Mn was maintained in the culture following viral inoculation. This inhibitory pattern was consistent with the reduction in HIV-1 infection observed in Figure 1B, further supporting the conclusion that Mn-mediated restriction occurs predominantly at the reverse transcription stage.

Having identified reverse transcription as a major step affected by Mn treatment, we next investigated whether Mn directly inhibits HIV-1 RT enzymatic activity. To address this possibility, we performed a cell-free HIV-1 RT activity assay. Since Mn exhibited potent antiviral activity in primary macrophages at concentrations of 25–50 µM, we first evaluated its direct effects on HIV-1 RT activity at these concentrations. As shown in Figure 3C, neither 25 nor 50 µM Mn significantly inhibited HIV-1 RT enzymatic activity.

We subsequently extended the analysis to a broader concentration range to determine whether higher Mn concentrations could affect RT activity. Consistent with our initial findings, Mn showed no inhibitory effects at concentrations up to 100 µM. However, at concentrations exceeding 100 µM, Mn inhibited HIV-1 RT activity in a concentration-dependent manner (Figure 3D). Notably, these inhibitory concentrations were substantially higher than those required to suppress HIV-1 infection in primary macrophages.

Those data indicate that the inhibition of HIV-1 reverse transcription observed in Mn-treated macrophages is unlikely to result from direct inhibition of viral RT enzymatic activity at the antiviral concentrations tested, suggesting the involvement of host cell-dependent mechanisms.

### Mn-Induced antiviral gene expression and the potential involvement of ATM signaling in HIV-1 restriction

Our previous studies demonstrated that Mn enhances innate antiviral signaling through ATM-TBK1 phosphorylation and regulates the expression of multiple host genes associated with antiviral defense [23]. To further investigate the host responses potentially contributing to Mn-mediated HIV-1 restriction, we examined the expression profiles of 10 selected Mn-responsive genes in primary human macrophages following Mn treatment for 1, 2, or 3 days. As shown in Figure 4A, Mn treatment induced distinct temporal patterns of gene expression. TNFAIP6, IL1A, CCL4, and CCL5 exhibited increased relative expression at later time points, whereas PLOD2 and ANKRD1 showed elevated expression at earlier time points. These findings suggest that Mn induces dynamic changes in host genes associated with innate immune responses and antiviral defense.

**Fig 4.**
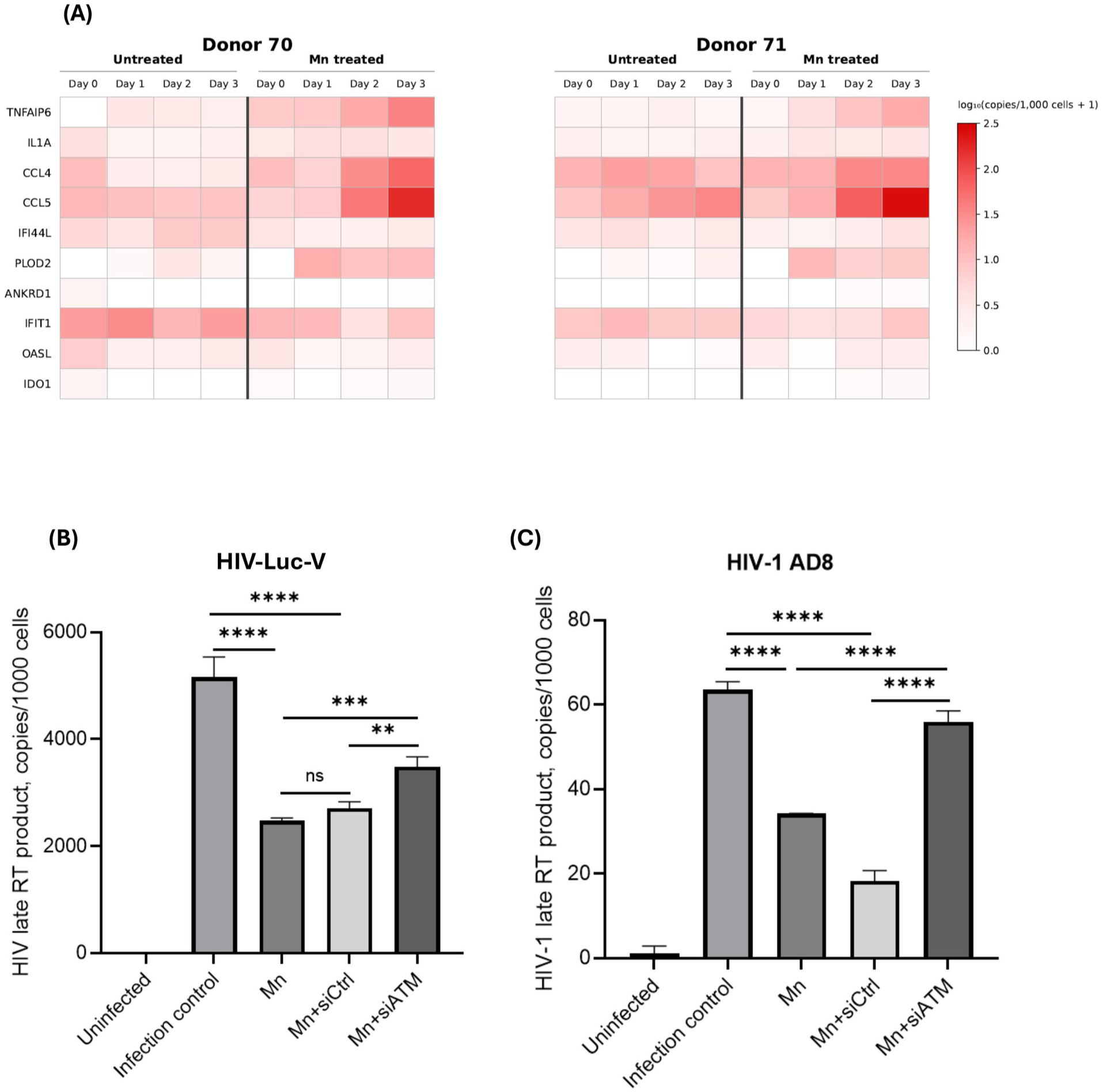
Mn-induced antiviral gene expression and the potential involvement of ATM signaling in HIV-1 restriction in human macrophages. **(A)** Heatmaps showing the temporal expression profiles of 10 selected Mn-responsive genes in MDMs treated with or without Mn for 1, 2, or 3 days. Gene expression was quantified by digital PCR, normalized to GAPDH, and expressed as copies per 1,000 cells. Heatmaps were generated using log_10_-transformed copy numbers [log_10_(copies/1,000 cells + 1)], with a shared color scale ranging from light to dark red indicating increasing expression levels. Data from two independent donors are shown. **(B, C)** Evaluation of ATM involvement in Mn-mediated HIV-1 restriction. MDMs were transfected with ATM-targeting siRNA (siATM) or non-targeting control siRNA (siCtrl), followed by Mn treatment and HIV-Luc-V **(B)** or HIV-1 AD8 **(C)** infection. HIV-1 late RT products were quantified by digital PCR and normalized to cellular RNase P copy numbers. Results are expressed as HIV-1 late RT copies per 1,000 cells. Both panels represent data from one of three independent donors. Error bars represent mean ± SD of technical replicates (n = 3). Statistical significance was determined by one-way ANOVA with Dunnett’s multiple comparisons test: ns, not significant; ** p < 0.01; *** p < 0.001; **** p < 0.0001.

To investigate the potential involvement of ATM signaling in Mn-mediated HIV-1 restriction, we examined the effects of ATM-targeting siRNA on HIV-1 reverse transcription in primary macrophages. Mn treatment markedly reduced HIV-1 late reverse transcription (RT) products compared with the untreated infection control (Fig. 4B). While non-targeting control siRNA did not significantly affect Mn-mediated inhibition, ATM-targeting siRNA partially restored HIV-1 late RT products in Mn-treated cells. Furthermore, Mn treatment similarly reduced the accumulation of HIV-1 late RT products following infection with replication-competent HIV-1 AD8 (Fig. 4C). Together, these findings further support reverse transcription as a major step targeted by Mn and provide preliminary evidence that ATM-associated signaling may contribute to Mn-mediated HIV-1 restriction in primary macrophages.

Overall, these findings indicate that Mn modulates the expression of multiple host immune-associated genes and provide preliminary evidence supporting a potential contribution of ATM-associated signaling to its antiviral activity.

## Discussion

In this study, we demonstrate that manganese (Mn) exerts potent, dose-dependent anti-HIV-1 activity in primary human macrophages, achieving up to 80%–90% viral suppression at a well-tolerated concentration of 20 or 25 µM. Importantly, Mn was effective both as a pretreatment and when administered post-infection, indicating that it can rapidly establish and maintain an antiviral state even after viral entry has been initiated. By systematically dissecting the viral replication cycle and comparing primary human immune subsets, our findings provide new mechanistic insights into how trace metal signaling interfaces with retroviral restriction.

A central finding of our work is that Mn inhibits HIV-1 replication specifically at the stage of reverse transcription. Further dissection revealed that Mn fails to directly inhibit recombinant HIV-1 reverse transcriptase (RT) activity in cell-free enzymatic assays at concentrations of 25 µM or 50 µM, despite these concentrations demonstrating potent anti-HIV activity in cells. However, Mn does directly inhibit HIV-1 RT enzymatic activity at much higher concentrations (100–5,000 µM). We also examined whether Mn interferes with other reverse transcriptases, such as MultiScribe RT, which is widely used for cDNA synthesis. As shown in S2 Fig, in the presence of 5.5 mM MgCl_2_, Mn dose-dependently inhibited MultiScribe RT activity, reducing cDNA synthesis of *GAPDH* and *CDKN1A*. Completely replacing MgCl_2_ with MnCl_2_ caused enzymatic activity to drop to 2.23% and 6.03%, respectively. Early biochemical studies suggested that supraphysiological levels of divalent Mn compete with magnesium (Mg) cofactors at the polymerase active site to impair retroviral DNA synthesis in vitro [39]. Our findings at higher concentrations are consistent with this previous report. However, our results indicate that at 25 µM or 50 µM in primary human macrophages, viral suppression is not driven by direct physical blockade of the viral enzyme. Instead, the dissociation between intracellular reverse transcription arrest and preserved cell-free RT activity strongly points toward an indirect, host-mediated restriction mechanism.

Consistent with the model proposed above, our transcriptomic analyses revealed that Mn induces a robust signature of innate antiviral defense and immunoregulatory genes. Notably, the antiviral activity of Mn was strictly cell-type-specific, occurring in primary macrophages while sparing primary dendritic cells and CD4+ T cells. Several biological factors may account for this selective restriction. First, primary human macrophages possess a distinct intracellular environment characterized by limited dNTP pools and high constitutive expression of myeloid-enriched restriction factors, such as SAMHD1, the APOBEC3 family, and MX2, which have been shown to tightly regulate reverse transcription and early post-entry steps [40–42]. Second, myeloid cells differ markedly from lymphocytes in their expression of divalent metal transporters (such as ZIP8, ZIP14, and DMT1) and innate nucleic acid sensors [15, 43, 44]. While macrophages mount vigorous type I interferon and interferon-stimulated gene (ISG) responses upon Mn-mediated sensitization of cGAS–STING or ATM–TBK1 pathways [19, 21, 23], resting or activated CD4+ T cells typically exhibit attenuated cytosolic DNA-sensing capacity [45, 46]. Similarly, dendritic cells may require distinct co-stimulatory thresholds or compartmentalized metal trafficking to trigger an equivalent antiviral state [47–49].

Given the potential involvement of host restriction factors, we first examined the role of SAMHD1, as its dephosphorylated form is specifically responsible for restricting the reverse transcription step of HIV-1 replication [50, 51]. However, Western blot analysis revealed no substantial difference in SAMHD1 phosphorylation regardless of whether the cells were treated with Mn (S3 Fig). Therefore, SAMHD1 is unlikely to be the primary driver of Mn-mediated anti-HIV activity.

Our previous studies demonstrated that Mn enhances innate antiviral signaling through the phosphorylation of multiple kinases, including ATM and TBK1 [21, 23]. To further delineate the role of innate immune pathways and assess the specific contribution of ATM, we analyzed host gene expression and performed knockdown assays. In the present study, Mn induced the expression of multiple host genes associated with antiviral defense, while preliminary ATM silencing experiments partially attenuated Mn-mediated HIV-1 restriction. These findings are consistent with a potential contribution of ATM-associated signaling to the antiviral effects of Mn. However, the precise mechanistic relationship between ATM, TBK1 activation, and the induction of antiviral effector genes remains to be established. Given the broad immunomodulatory effects of Mn, multiple host pathways likely act cooperatively to restrict HIV-1 reverse transcription in macrophages.

Macrophages play a critical role in HIV-1 pathogenesis as long-lived, tissue-resident cells that disseminate the virus to anatomical sanctuaries including the central nervous system, lymph nodes, and gut-associated lymphoid tissue, and contribute to persistent viral reservoirs during antiretroviral therapy (ART) [52–55]. Identifying host-directed agents capable of selectively fortifying macrophages against HIV-1 replication is therefore of substantial therapeutic interest. Because Mn acts through host innate signaling networks rather than targeting mutable viral proteins, Mn-based strategies could offer a high genetic barrier against drug resistance and complement existing antiretroviral regimens [56–58].

Several limitations of our study warrant consideration and future investigation. First, while our transcriptomic data implicate host innate defense pathways in blocking reverse transcription, targeted loss-of-function studies, such as siRNA-mediated knockdown or CRISPR-Cas9 knockout of specific ISGs, will be required to pinpoint the exact downstream effectors indispensable for Mn-mediated restriction. Second, we did not quantify intracellular Mn accumulation across macrophages, dendritic cells, and CD4+ T cells; differential metal uptake or subcellular sequestration could contribute to the observed cell-type specificity. Finally, systemic administration of free Mn salts in vivo is constrained by rapid clearance and potential neurotoxicity at high doses [59, 60]. Future translational efforts should therefore evaluate targeted delivery platforms, such as macrophage-tropic nanoparticles or colloidal manganese formulations to safely harness Mn-driven innate immunity in vivo [61, 62].

In summary, our work demonstrates that Mn restricts HIV-1 in primary human macrophages by triggering a host-directed antiviral program that arrests reverse transcription without directly inhibiting the viral reverse transcriptase. These findings deepen our understanding of metalloimmunology in retroviral infection and highlight macrophage-targeted immunomodulation as a promising avenue for future anti-HIV-1 strategies.

## Acknowledgments

The authors sincerely thank Dr. Veronique Nussenblatt for her support of this project. The following reagent was obtained through BEI Resources, NIAID, NIH: HIV-1 NL4-3 ΔEnv luciferase reporter plasmid (HRP-3418).

## Author Contributions

**Conceptualization:** HS. **Data curation:** HS. **Formal analysis:** HS. **Funding acquisition:** TI. **Investigation:** HS, RWR, WB, JM. **Methodology:** HS. **Project administration:** HS, TI. **Resources:** TI. **Supervision:** HS, AR, SL, TI. **Visualization:** HS, RW-R, WB. **Writing – original draft:** HS. **Writing – review & editing:** HS, RW-R, AR, SL, TI.

## Data Availability

All relevant data are within the manuscript and its Supporting Information files.

## Supporting Information

**S1 Fig.**
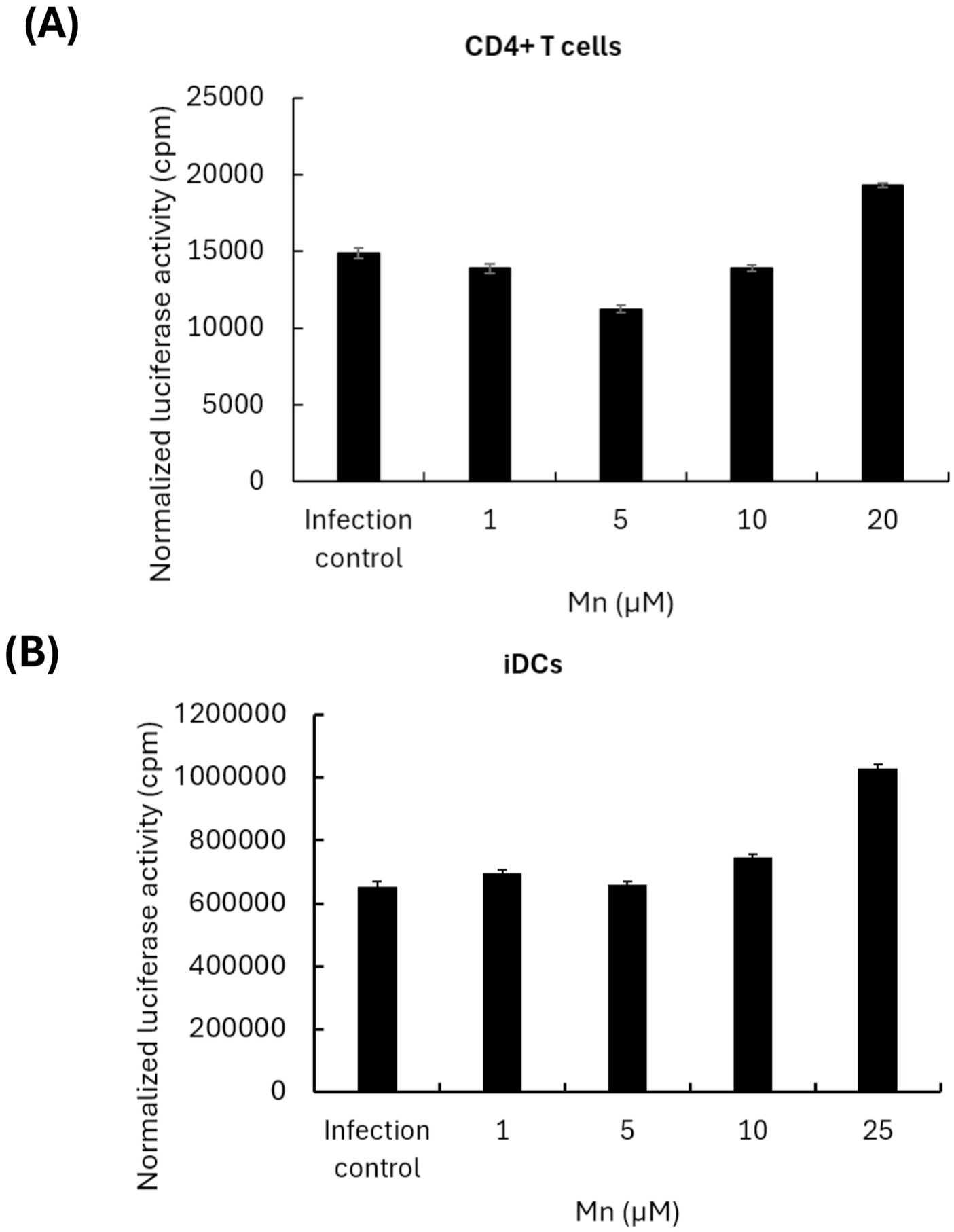
Mn pretreatment does not inhibit HIV-1 infection in primary CD4+ T cells and immature dendritic cells. Primary human CD4+ T cells (A) and monocyte-derived immature dendritic cells (iDCs) (B) were pretreated with the indicated concentrations of Mn for 48 h before infection with VSV-G-pseudotyped HIV-1 luciferase reporter virus (HIV-Luc-V HIV-1 infection was assessed by luciferase activity normalized by total protein concentration for each sample. Data are presented as mean ± SD (*n* = 3) and are representative of three independent donors.

**S2 Fig.**
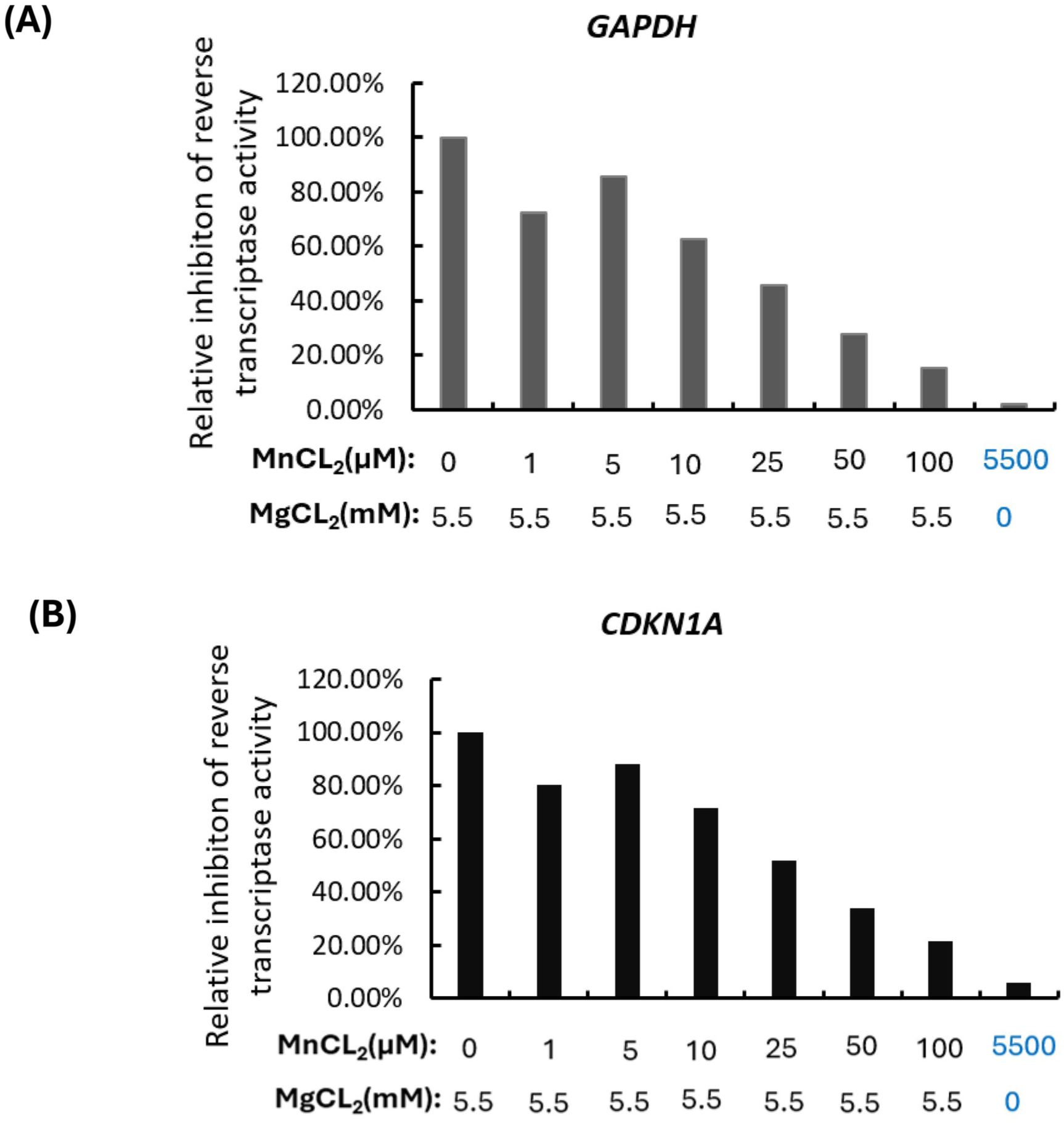
Mn inhibits MultiScribe reverse transcriptase activity in a concentration-dependent manner. The effects of Mn on MultiScribe reverse transcriptase (RT) activity were evaluated by adding increasing concentrations of MnCl_2_ to cDNA synthesis reactions containing 5.5 mM MgCl_2_ or by completely replacing 5.5 mM MgCl_2_ with an equivalent concentration of MnCl_2_ (5.5 mM). The resulting cDNA copy numbers of *GAPDH* **(A)** and *CDKN1A* **(B)** were quantified by digital PCR. RT activity was normalized to that of the standard reaction containing 5.5 mM MgCl_2_ and expressed as a percentage relative to the corresponding control (*n* = 3).

**S3 Fig.**
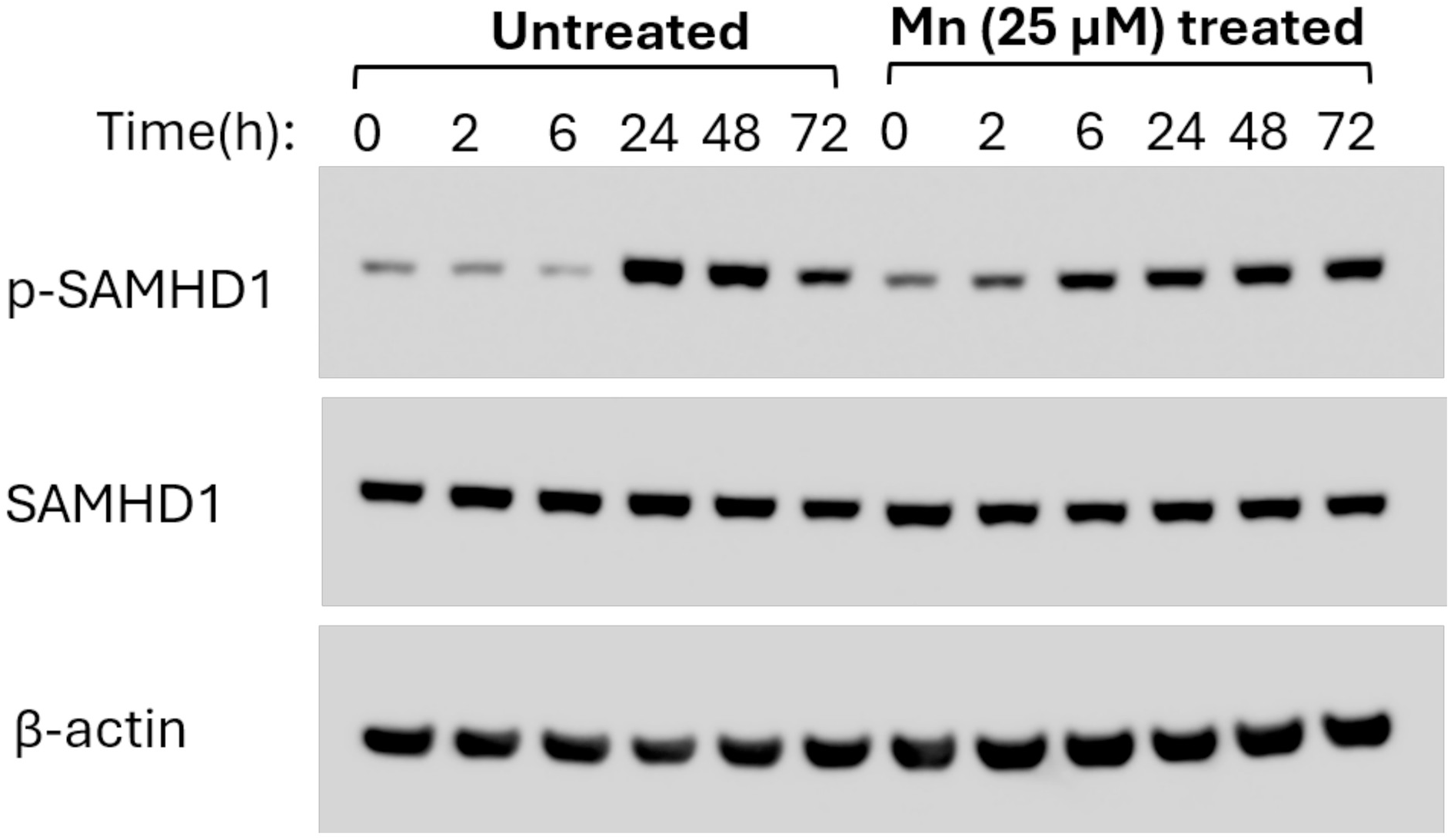
Mn treatment does not induce SAMHD1 dephosphorylation in primary human macrophages. Primary human monocyte-derived macrophages (MDMs) were treated with 25 µM Mn for the indicated time periods. Whole-cell lysates were collected and subjected to Western blot analysis using antibodies against phosphorylated SAMHD1 (p-SAMHD1), total SAMHD1, and β-actin. Representatives immunoblot results from one of three independent donors are shown.

## References

1. Carter A, Zhang M, Trejo BG, Verma M, Brewer ED, Lasher D, et al. Global, regional, and national burden of HIV/AIDS, 1990–2023, with the estimated burden attributable to intimate partner violence and forecasted impacts of funding cuts through 2030: results from the Global Burden of Disease Study 2023. The Lancet HIV. 2026. doi: 10.1016/S2352-3018(26)00147-5.

2. HIV/AIDS JUNPo. Ending AIDS starts with access: World AIDS Day report 2026. Geneva: UNAIDS, 2026.

3. Pace MJ, Agosto L, Graf EH, O’Doherty U. HIV reservoirs and latency models. Virology. 2011;411(2):344–54. doi: 10.1016/j.virol.2010.12.041.

4. Siliciano JD, Siliciano RF. In Vivo Dynamics of the Latent Reservoir for HIV-1: New Insights and Implications for Cure. Annual Review of Pathology: Mechanisms of Disease. 2022;17(Volume 17, 2022):271–94. doi: 10.1146/annurev-pathol-050520-112001.

5. Li JZ, Aga E, Bosch RJ, Pilkinton M, Kroon E, MacLaren L, et al. Time to Viral Rebound After Interruption of Modern Antiretroviral Therapies. Clin Infect Dis. 2022;74(5):865–70. doi: 10.1093/cid/ciab541. PubMed PMID: 34117753; PubMed Central PMCID: PMCPMC8906742.

6. Gunst JD, Gohil J, Li JZ, Bosch RJ, White CSA, Chun T-W, et al. Time to HIV viral rebound and frequency of post-treatment control after analytical interruption of antiretroviral therapy: an individual data-based meta-analysis of 24 prospective studies. Nature Communications. 2025;16(1):906. doi: 10.1038/s41467-025-56116-1.

7. Chawla A, Wang C, Patton C, Murray M, Punekar Y, de Ruiter A, et al. A Review of Long-Term Toxicity of Antiretroviral Treatment Regimens and Implications for an Aging Population. Infect Dis Ther. 2018;7(2):183–95. Epub 20180514. doi: 10.1007/s40121-018-0201-6. PubMed PMID: 29761330; PubMed Central PMCID: PMCPMC5986685.

8. Flavell AJ. Retroelements, reverse transcriptase and evolution. Comp Biochem Physiol B Biochem Mol Biol. 1995;110(1):3–15. doi: 10.1016/0305-0491(94)00122-b. PubMed PMID: 7532085.

9. Coffin JM HS, Varmus HE, editor. Retroviruses. Cold Spring Harbor, NY: Cold Spring Harbor Laboratory Press; 1997.

10. Telesnitsky A, Goff SP. Reverse Transcriptase and the Generation of Retroviral DNA. In: Coffin JM, Hughes SH, Varmus HE, editors. Retroviruses. Cold Spring Harbor (NY): Cold Spring Harbor Laboratory Press Copyright © 1997, Cold Spring Harbor Laboratory Press.; 1997.

11. Sarafianos SG, Marchand B, Das K, Himmel DM, Parniak MA, Hughes SH, et al. Structure and function of HIV-1 reverse transcriptase: molecular mechanisms of polymerization and inhibition. J Mol Biol. 2009;385(3):693–713. Epub 20081103. doi: 10.1016/j.jmb.2008.10.071. PubMed PMID: 19022262; PubMed Central PMCID: PMCPMC2881421.

12. Arts EJ, Hazuda DJ. HIV-1 antiretroviral drug therapy. Cold Spring Harb Perspect Med. 2012;2(4):a007161. doi: 10.1101/cshperspect.a007161. PubMed PMID: 22474613; PubMed Central PMCID: PMCPMC3312400.

13. Clutter DS, Jordan MR, Bertagnolio S, Shafer RW. HIV-1 drug resistance and resistance testing. Infect Genet Evol. 2016;46:292–307. Epub 20160829. doi: 10.1016/j.meegid.2016.08.031. PubMed PMID: 27587334; PubMed Central PMCID: PMCPMC5136505.

14. Wang C, Zhang R, Wei X, Lv M, Jiang Z. Metalloimmunology: The metal ion-controlled immunity. Adv Immunol. 2020;145:187–241. Epub 20191209. doi: 10.1016/bs.ai.2019.11.007. PubMed PMID: 32081198.

15. Murdoch CC, Skaar EP. Nutritional immunity: the battle for nutrient metals at the host-pathogen interface. Nat Rev Microbiol. 2022;20(11):657–70. Epub 20220531. doi: 10.1038/s41579-022-00745-6. PubMed PMID: 35641670; PubMed Central PMCID: PMCPMC9153222.

16. Sun X, Zhou X, Shi X, Abed OA, An X, Lei YL, et al. Strategies for the development of metalloimmunotherapies. Nat Biomed Eng. 2024;8(9):1073–91. Epub 20240624. doi: 10.1038/s41551-024-01221-7. PubMed PMID: 38914800; PubMed Central PMCID: PMCPMC11410547.

17. Li L, Yang X. The Essential Element Manganese, Oxidative Stress, and Metabolic Diseases: Links and Interactions. Oxid Med Cell Longev. 2018;2018:7580707. Epub 20180405. doi: 10.1155/2018/7580707. PubMed PMID: 29849912; PubMed Central PMCID: PMCPMC5907490.

18. Chen P, Bornhorst J, Aschner M. Manganese metabolism in humans. Front Biosci (Landmark Ed). 2018;23(9):1655–79. Epub 20180301. doi: 10.2741/4665. PubMed PMID: 29293455.

19. Wang C, Guan Y, Lv M, Zhang R, Guo Z, Wei X, et al. Manganese Increases the Sensitivity of the cGAS-STING Pathway for Double-Stranded DNA and Is Required for the Host Defense against DNA Viruses. Immunity. 2018;48(4):675–87.e7. doi: 10.1016/j.immuni.2018.03.017.

20. Zhao Z, Ma Z, Wang B, Guan Y, Su X-D, Jiang Z. Mn2+ Directly Activates cGAS and Structural Analysis Suggests Mn2+ Induces a Noncanonical Catalytic Synthesis of 2′3′-cGAMP. Cell Reports. 2020;32(7):108053. doi: 10.1016/j.celrep.2020.108053.

21. Sui H, Chen Q, Yang J, Srirattanapirom S, Imamichi T. Manganese enhances DNA- or RNA-mediated innate immune response by inducing phosphorylation of TANK-binding kinase 1. iScience. 2022;25(11):105352. Epub 20221013. doi: 10.1016/j.isci.2022.105352. PubMed PMID: 36325059; PubMed Central PMCID: PMCPMC9619380.

22. Sun S, Xu Y, Qiu M, Jiang S, Cao Q, Luo J, et al. Manganese Mediates Its Antiviral Functions in a cGAS-STING Pathway Independent Manner. Viruses. 2023;15(3). Epub 20230228. doi: 10.3390/v15030646. PubMed PMID: 36992355; PubMed Central PMCID: PMCPMC10058264.

23. Sui H, Wiscovitch-Russo R, Cachaco S, Yang J, Bruchey W, Laverdure S, et al. Manganese mediates antiviral effects by driving an ATM - TBK1 phosphorylation signaling pathway. Frontiers in Immunology. 2025;Volume 16-2025. doi: 10.3389/fimmu.2025.1708516.

24. Yu L, Chang H, Xie W, Zheng Y, Yang L, Wu Q, et al. Manganese is a potent inducer of lysosomal activity that inhibits de novo HBV infection. PLOS Pathogens. 2025;21(1):e1012800. doi: 10.1371/journal.ppat.1012800.

25. Zhang Y, Han S, Li Y, Zhou Y, Sun M, Hu M, et al. Manganese inhibits HBV transcription and promotes HBsAg degradation at non-toxic levels. Int J Biol Macromol. 2024;280(Pt 2):135764. Epub 20240917. doi: 10.1016/j.ijbiomac.2024.135764. PubMed PMID: 39299429.

26. Shi PD, Xu YP, Zhu Z, Zhou C, Wu M, He Y, et al. Manganese Mineralization of Pathogenic Viruses as a Universal Vaccine Platform. Adv Sci (Weinh). 2023;10(33):e2303615. Epub 20231022. doi: 10.1002/advs.202303615. PubMed PMID: 37867242; PubMed Central PMCID: PMCPMC10667830.

27. Zhang Z, Zhang R, Yin J, Zhao S, Qin X, Chen F, et al. Antiviral Effect of Manganese against Foot-and-Mouth Disease Virus Both in PK15 Cells and Mice. Viruses. 2023;15(2):390. PubMed PMID: 10.3390/v15020390.

28. Lahaye X, Satoh T, Gentili M, Cerboni S, Conrad C, Hurbain I, et al. The capsids of HIV-1 and HIV-2 determine immune detection of the viral cDNA by the innate sensor cGAS in dendritic cells. Immunity. 2013;39(6):1132–42. Epub 20131121. doi: 10.1016/j.immuni.2013.11.002. PubMed PMID: 24269171.

29. Rasaiyaah J, Tan CP, Fletcher AJ, Price AJ, Blondeau C, Hilditch L, et al. HIV-1 evades innate immune recognition through specific cofactor recruitment. Nature. 2013;503(7476):402–5. Epub 20131106. doi: 10.1038/nature12769. PubMed PMID: 24196705; PubMed Central PMCID: PMCPMC3928559.

30. Theodore TS, Englund G, Buckler-White A, Buckler CE, Martin MA, Peden KW. Construction and characterization of a stable full-length macrophage-tropic HIV type 1 molecular clone that directs the production of high titers of progeny virions. AIDS Res Hum Retroviruses. 1996;12(3):191–4. doi: 10.1089/aid.1996.12.191. PubMed PMID: 8835195.

31. Chang LJ, Urlacher V, Iwakuma T, Cui Y, Zucali J. Efficacy and safety analyses of a recombinant human immunodeficiency virus type 1 derived vector system. Gene Ther. 1999;6(5):715–28. doi: 10.1038/sj.gt.3300895. PubMed PMID: 10505094.

32. Connor RI, Chen BK, Choe S, Landau NR. Vpr is required for efficient replication of human immunodeficiency virus type-1 in mononuclear phagocytes. Virology. 1995;206(2):935–44. doi: 10.1006/viro.1995.1016. PubMed PMID: 7531918.

33. He J, Choe S, Walker R, Di Marzio P, Morgan DO, Landau NR. Human immunodeficiency virus type 1 viral protein R (Vpr) arrests cells in the G2 phase of the cell cycle by inhibiting p34cdc2 activity. J Virol. 1995;69(11):6705–11. doi: 10.1128/jvi.69.11.6705-6711.1995. PubMed PMID: 7474080; PubMed Central PMCID: PMCPMC189580.

34. Schaeffer E, Soros VB, Greene WC. Compensatory link between fusion and endocytosis of human immunodeficiency virus type 1 in human CD4 T lymphocytes. J Virol. 2004;78(3):1375–83. doi: 10.1128/jvi.78.3.1375-1383.2004. PubMed PMID: 14722292; PubMed Central PMCID: PMCPMC321368.

35. Imamichi T, Chen Q, Sowrirajan B, Yang J, Laverdure S, Marquez M, et al. Interleukin-27-induced HIV-resistant dendritic cells suppress reveres transcription following virus entry in an SPTBN1, autophagy, and YB-1 independent manner. PLOS ONE. 2023;18(11):e0287829. doi: 10.1371/journal.pone.0287829.

36. Dai L, Lidie KB, Chen Q, Adelsberger JW, Zheng X, Huang D, et al. IL-27 inhibits HIV-1 infection in human macrophages by down-regulating host factor SPTBN1 during monocyte to macrophage differentiation. J Exp Med. 2013;210(3):517–34. Epub 20130304. doi: 10.1084/jem.20120572. PubMed PMID: 23460728; PubMed Central PMCID: PMCPMC3600911.

37. Cavrois M, de Noronha C, Greene WC. A sensitive and specific enzyme-based assay detecting HIV-1 virion fusion in primary T lymphocytes. Nature Biotechnology. 2002;20(11):1151–4. doi: 10.1038/nbt745.

38. Butler SL, Hansen MS, Bushman FD. A quantitative assay for HIV DNA integration in vivo. Nat Med. 2001;7(5):631–4. doi: 10.1038/87979. PubMed PMID: 11329067.

39. Bolton EC, Mildvan AS, Boeke JD. Inhibition of Reverse Transcription In Vivo by Elevated Manganese Ion Concentration. Molecular Cell. 2002;9(4):879–89. doi: 10.1016/S1097-2765(02)00495-1.

40. Laguette N, Sobhian B, Casartelli N, Ringeard M, Chable-Bessia C, Ségéral E, et al. SAMHD1 is the dendritic-and myeloid-cell-specific HIV-1 restriction factor counteracted by Vpx. Nature. 2011;474(7353):654–7. doi: 10.1038/nature10117.

41. Malim MH, Bieniasz PD. HIV Restriction Factors and Mechanisms of Evasion. Cold Spring Harbor perspectives in medicine. 2012;2 5:a006940.

42. Goujon C, Moncorgé O, Bauby H, Doyle T, Ward CC, Schaller T, et al. Human MX2 is an interferon-induced post-entry inhibitor of HIV-1 infection. Nature. 2013;502(7472):559–62. Epub 20130918. doi: 10.1038/nature12542. PubMed PMID: 24048477; PubMed Central PMCID: PMCPMC3808269.

43. Sapkota M, Knoell DL. Essential Role of Zinc and Zinc Transporters in Myeloid Cell Function and Host Defense against Infection. J Immunol Res. 2018;2018:4315140. Epub 20181017. doi: 10.1155/2018/4315140. PubMed PMID: 30417019; PubMed Central PMCID: PMCPMC6207864.

44. Beker Aydemir T, Chang S-M, Guthrie GJ, Maki AB, Ryu M-S, Karabiyik A, et al. Zinc Transporter ZIP14 Functions in Hepatic Zinc, Iron and Glucose Homeostasis during the Innate Immune Response (Endotoxemia). PLOS ONE. 2012;7(10):e48679. doi: 10.1371/journal.pone.0048679.

45. Vermeire J, Roesch F, Sauter D, Rua R, Hotter D, Van Nuffel A, et al. HIV Triggers a cGAS-Dependent, Vpu- and Vpr-Regulated Type I Interferon Response in CD4+ T Cells. Cell Reports. 2016;17(2):413–24. doi: 10.1016/j.celrep.2016.09.023.

46. Elsner C, Ponnurangam A, Kazmierski J, Zillinger T, Jansen J, Todt D, et al. Absence of cGAS-mediated type I IFN responses in HIV-1-infected T cells. Proc Natl Acad Sci U S A. 2020;117(32):19475–86. Epub 20200724. doi: 10.1073/pnas.2002481117. PubMed PMID: 32709741; PubMed Central PMCID: PMCPMC7431009.

47. Rodriguez Rl, Müller S, Colombeau L, Solier Sp, Sindikubwabo F, Cañeque T. Metal Ion Signaling in Biomedicine. Chemical Reviews. 2025;125(2):660–744. doi: 10.1021/acs.chemrev.4c00577.

48. Cheng H, Chen W, Lin Y, Zhang J, Song X, Zhang D. Signaling pathways involved in the biological functions of dendritic cells and their implications for disease treatment. Mol Biomed. 2023;4(1):15. Epub 20230515. doi: 10.1186/s43556-023-00125-3. PubMed PMID: 37183207; PubMed Central PMCID: PMCPMC10183318.

49. Wang Z, Chen Q, Chen X, Lv L, Liu W. New exploration of the effect of intracellular metal ions on tumor immunity. Coordination Chemistry Reviews. 2026;555:217594. doi: 10.1016/j.ccr.2026.217594.

50. White Tommy E, Brandariz-Nuñez A, Valle-Casuso Jose C, Amie S, Nguyen Laura A, Kim B, et al. The Retroviral Restriction Ability of SAMHD1, but Not Its Deoxynucleotide Triphosphohydrolase Activity, Is Regulated by Phosphorylation. Cell Host & Microbe. 2013;13(4):441–51. doi: 10.1016/j.chom.2013.03.005.

51. Cribier A, Descours B, Valadão Ana Luiza C, Laguette N, Benkirane M. Phosphorylation of SAMHD1 by Cyclin A2/CDK1 Regulates Its Restriction Activity toward HIV-1. Cell Reports. 2013;3(4):1036–43. doi: 10.1016/j.celrep.2013.03.017.

52. Kruize Z, Kootstra NA. The Role of Macrophages in HIV-1 Persistence and Pathogenesis. Frontiers in Microbiology. 2019;Volume 10 - 2019. doi: 10.3389/fmicb.2019.02828.

53. Ganor Y, Real F, Sennepin A, Dutertre CA, Prevedel L, Xu L, et al. HIV-1 reservoirs in urethral macrophages of patients under suppressive antiretroviral therapy. Nat Microbiol. 2019;4(4):633–44. Epub 20190204. doi: 10.1038/s41564-018-0335-z. PubMed PMID: 30718846.

54. Honeycutt JB, Thayer WO, Baker CE, Ribeiro RM, Lada SM, Cao Y, et al. HIV persistence in tissue macrophages of humanized myeloid-only mice during antiretroviral therapy. Nat Med. 2017;23(5):638–43. Epub 20170417. doi: 10.1038/nm.4319. PubMed PMID: 28414330; PubMed Central PMCID: PMCPMC5419854.

55. Veenhuis RT, Abreu CM, Costa PAG, Ferreira EA, Ratliff J, Pohlenz L, et al. Monocyte-derived macrophages contain persistent latent HIV reservoirs. Nature Microbiology. 2023;8(5):833–44. doi: 10.1038/s41564-023-01349-3.

56. Kaufmann SHE, Dorhoi A, Hotchkiss RS, Bartenschlager R. Host-directed therapies for bacterial and viral infections. Nat Rev Drug Discov. 2018;17(1):35–56. Epub 20170922. doi: 10.1038/nrd.2017.162. PubMed PMID: 28935918; PubMed Central PMCID: PMCPMC7097079.

57. Kumar N, Sharma S, Kumar R, Tripathi BN, Barua S, Ly H, et al. Host-Directed Antiviral Therapy. Clin Microbiol Rev. 2020;33(3). Epub 20200513. doi: 10.1128/cmr.00168-19. PubMed PMID: 32404434; PubMed Central PMCID: PMCPMC7227448.

58. Gu X, Zheng M, Gao Y, Lin S, Zhang X, Chen C, et al. Overview of host-directed antiviral targets for future research and drug development. Acta Pharmaceutica Sinica B. 2025;15(4):1723–51. doi: 10.1016/j.apsb.2025.03.011.

59. O’Neal SL, Zheng W. Manganese Toxicity Upon Overexposure: a Decade in Review. Curr Environ Health Rep. 2015;2(3):315–28. doi: 10.1007/s40572-015-0056-x. PubMed PMID: 26231508; PubMed Central PMCID: PMCPMC4545267.

60. Peres TV, Schettinger MR, Chen P, Carvalho F, Avila DS, Bowman AB, et al. “Manganese-induced neurotoxicity: a review of its behavioral consequences and neuroprotective strategies”. BMC Pharmacol Toxicol. 2016;17(1):57. Epub 20161104. doi: 10.1186/s40360-016-0099-0. PubMed PMID: 27814772; PubMed Central PMCID: PMCPMC5097420.

61. Zhang R, Wang C, Guan Y, Wei X, Sha M, Yi M, et al. Manganese salts function as potent adjuvants. Cell Mol Immunol. 2021;18(5):1222–34. Epub 20210325. doi: 10.1038/s41423-021-00669-w. PubMed PMID: 33767434; PubMed Central PMCID: PMCPMC8093200.

62. Sun Y, Yin Y, Gong L, Liang Z, Zhu C, Ren C, et al. Manganese nanodepot augments host immune response against coronavirus. Nano Res. 2021;14(5):1260–72. Epub 20201229. doi: 10.1007/s12274-020-3243-5. PubMed PMID: 33391623; PubMed Central PMCID: PMCPMC7770383.

